## Supplementary figures and images for "Long non-coding RNA expression levels modulate cell-type specific splicing patterns by altering their interaction landscape with RNA-binding proteins"

### Supplementary material 4.pdf

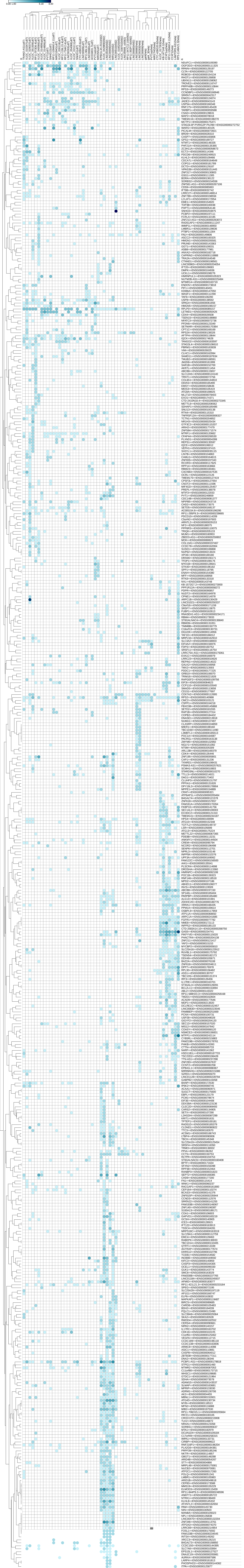

### Supplementary material 5.pdf

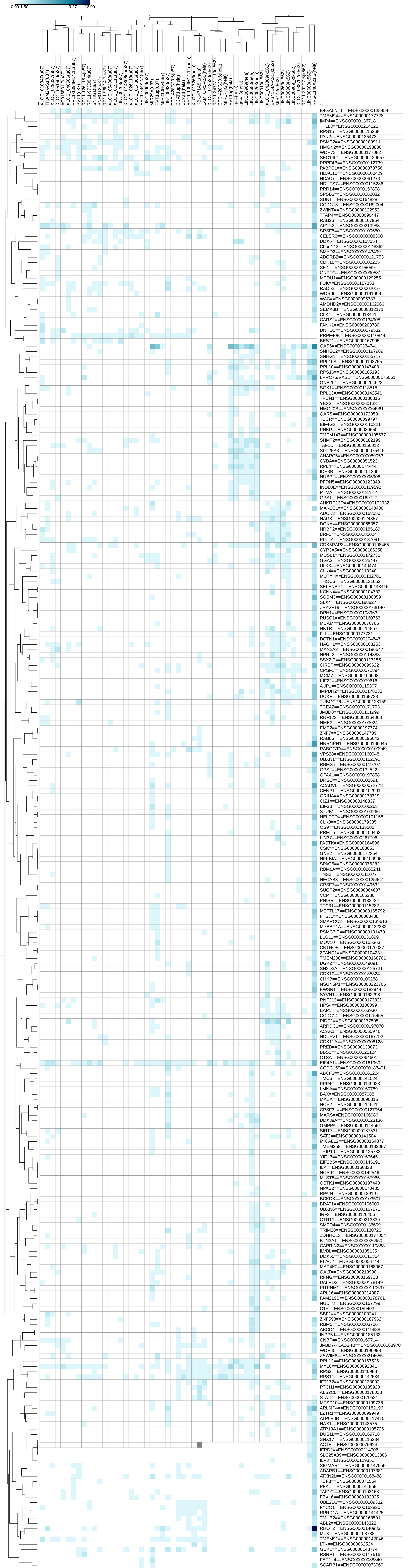

### Supplementary material 6.png

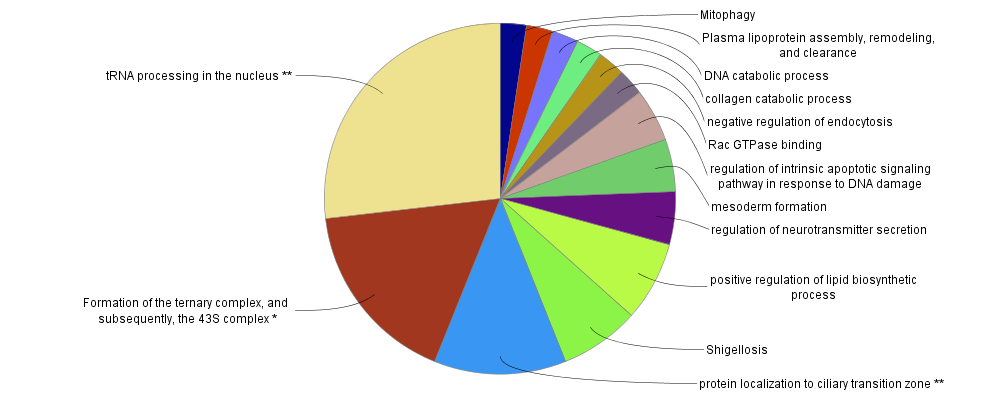

### Supplementary material 7.png

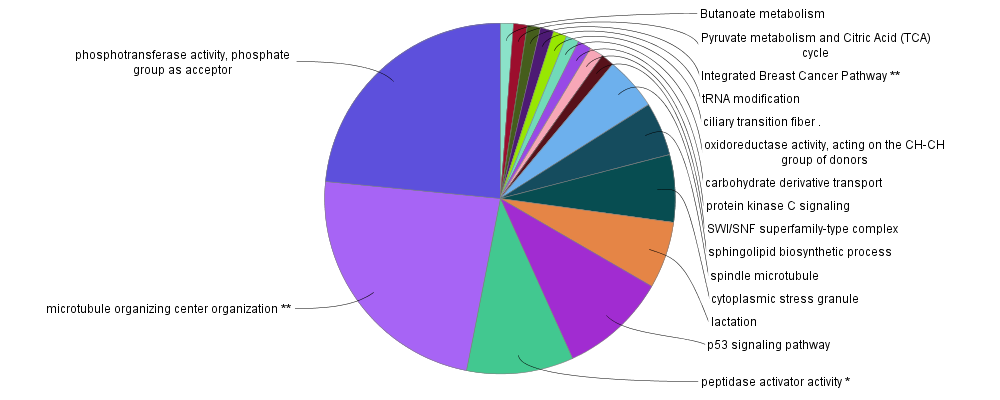

### Supplementary material 8.png

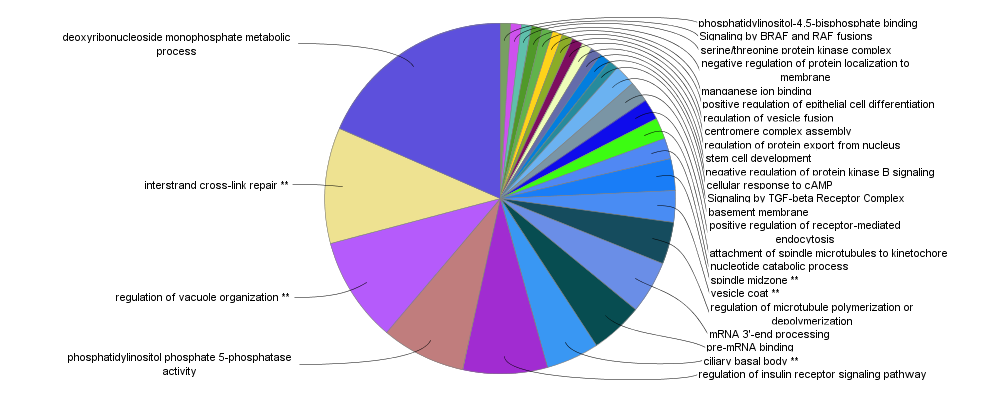

### Supplementary material 9.png

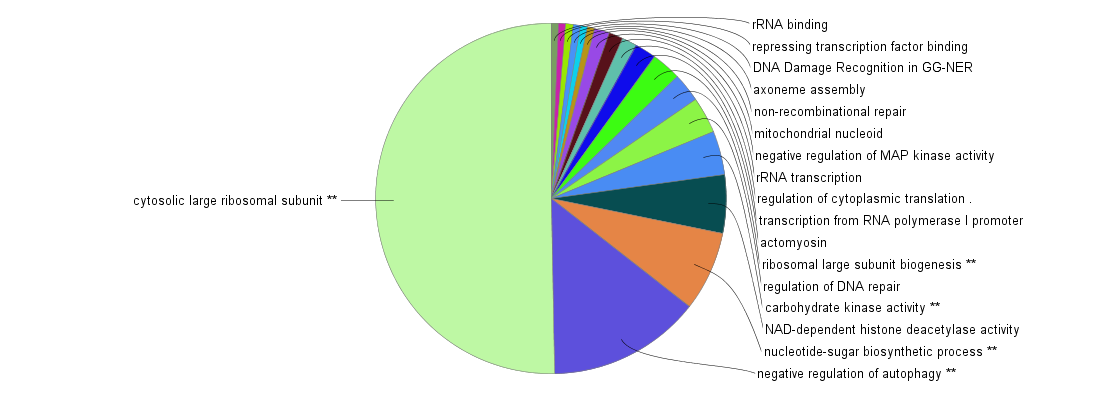

### Supplementary material 10.png

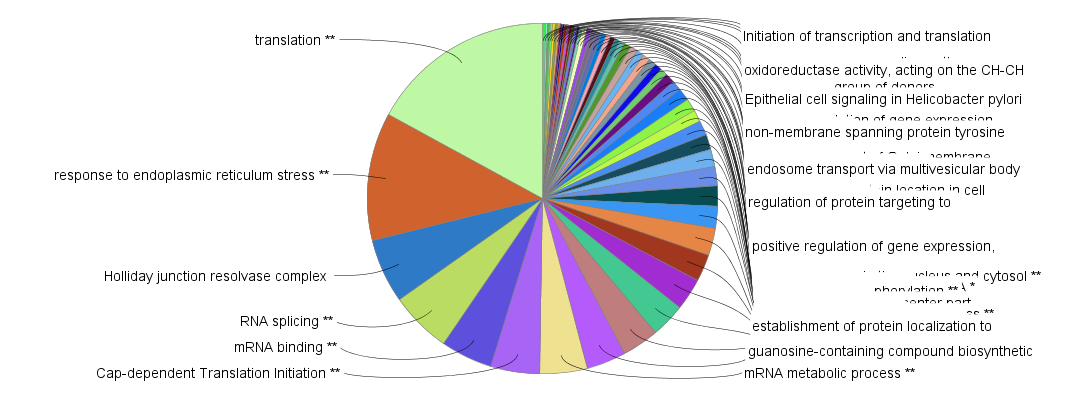
